# Spec2Class: Accurate Prediction of Plant Secondary Metabolite Class using Deep Learning

**DOI:** 10.1101/2024.03.17.585408

**Authors:** Victoria Poltorak, Nir Shachaf, Asaph Aharoni, David Zeevi

## Abstract

Mass spectrometry (MS)-based data is commonly used in studying metabolism and natural products, but typically requires domain-specific skill and experience to analyze. Existing computational tools for non-targeted metabolite analysis (i.e., metabolomics) mostly rely on comparison to reference MS spectral libraries for metabolite identification, limiting the annotation of metabolites for which reference spectra do not exist. This is the case in plant secondary metabolites, where most spectral features remain unidentified. Here, we developed *Spec2Class*, a deep-learning algorithm for the identification and classification of plant secondary metabolites from liquid chromatography (LC)-MS/MS spectra. We used the in-house spectral library of 7973 plant metabolite chemical standards, alongside publicly available data, to train *Spec2Class* to classify LC-MS/MS spectra to 43 common plant secondary metabolite classes. Tested on held out sets, our algorithm achieved an overall accuracy of 73%, outperforming state-of-the-art classification. We further established a prediction certainty parameter to set a threshold for low-confidence results. Applying this threshold, we reached an accuracy of 93% on an unseen dataset. We show a high robustness of our prediction to noise and to the data acquisition method. *Spec2Class* is publicly available and is anticipated to facilitate metabolite identification and accelerate natural product discovery.

**Significance Statement:** Untargeted mass spectrometry (MS) is essential for natural product discovery but is limited by product identification, which is often manual and requires domain-specific skills. *Spec2Class* addresses this limitation by accurately classifying plant secondary metabolites from LC-MS/MS spectra without reliance on reference spectral libraries. Trained on a substantial dataset and using a prediction certainty threshold, it outperforms state-of-the-art algorithms with 93% accuracy. This tool demonstrates high robustness against noise and different data acquisition methods, promising to streamline metabolite identification and expedite natural product research. *Spec2Class* is open-source, publicly available, and easy to integrate into natural product discovery pipelines.

## Introduction

The plant kingdom is estimated to produce nearly 1 million metabolites (1), of which only ∼200,000 are currently structurally determined and classified (2). A large portion of these metabolites are secondary (or specialized), small molecules which perform various functions, mainly as part of the numerous interactions between plants and other organisms (3).

One important tool for identifying plant metabolites is liquid chromatography coupled with mass spectrometry (LC-MS) which is particularly effective for rapidly determining the components of complex mixtures of molecules. Its high sensitivity and modularity allows it to be customized for a wide range of purposes such as reaction monitoring, reactant quantification, or support for de novo compound discovery (4). These advantages are often overcome with challenging data processing and interpretation, making MS-based metabolite annotations one of the major challenges in the field.

Analysis of metabolites is largely divided into targeted assays, in which a query consisting of a metabolite, or a set of metabolites is searched; and untargeted ones, in which an analysis of all mass features is carried out. Untargeted assays pose a bigger challenge, as they are unbiased and often not supported by previous knowledge. In such assays, metabolite identification is often conducted by matching the output mass spectra against reference spectra (5). To date and for the foreseeable future, these spectral libraries are incomplete and have a partial representation of many chemical classes.

This drawback gave rise to the development of computational tools that facilitate automated metabolite annotation by approaches such as library expansion with in-silico fragmentation predictions (6, 7) and molecular network analysis based on spectral similarity (8, 9) to other known compounds. Still, and despite these advances, only 2-5% of MS/MS spectra from experiments are fully annotated (10). Other tools in the field chose to aim for the metabolite’s chemical class association rather than full structure resolution. Albeit not as specific, metabolite chemical class assignment is very valuable for downstream analysis. For chemical assignment, a major leap in the field can be attributed to *CANOPUS (11)* and *SteroidXtract* (12) which used machine learning and, specifically, deep learning algorithms for the task.

Here we present *Spec2Class*, a machine learning classifier designed to facilitate plant secondary metabolite identification, through chemical class determination (Fig. 1a). We classify secondary metabolites to 43 common plant chemical superclasses, based on the NPClassifier ontology. We trained and tested our models on a unique in-house spectral library of 7,973 plant secondary metabolite standards, which we supplemented with additional 13,689 publicly available spectra from the Mass-bank of North America (MoNA; https://mona.fiehnlab.ucdavis.edu/). Our method predicts chemical classes from a mass spectra obtained via positive ionization mode data dependent acquisition (DDA) or data independent acquisition (DIA) in a high-resolution Quadrupole Time of Flight (QToF) MS instrument.

**Figure 1.**
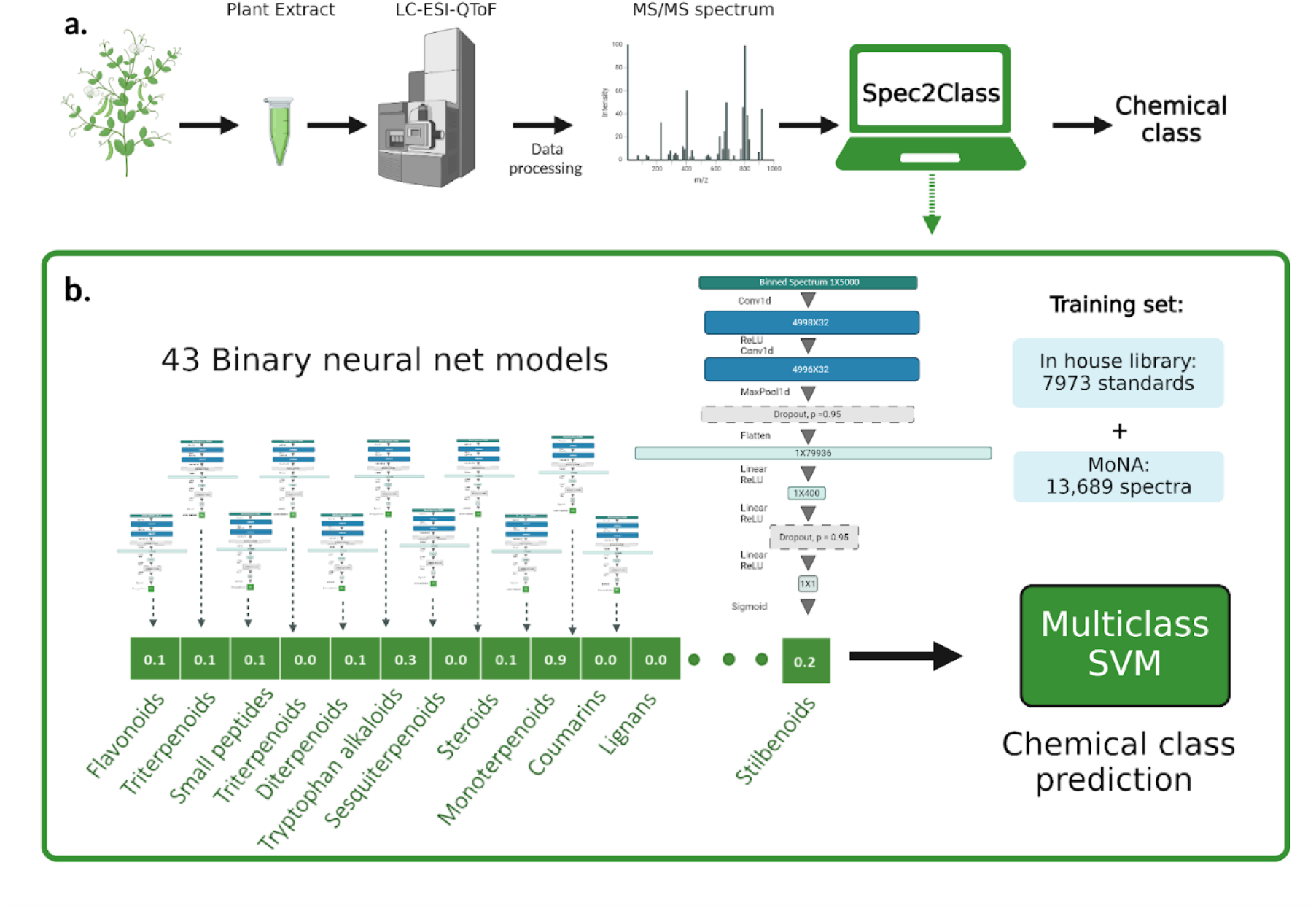
An overview of the *Spec2Class* pipeline. **(a)** A depiction of how *Spec2Class* fits into a mass spectrometry analysis pipeline. **(b)** *Spec2Class* has three main steps: (1) spectra parsing and binning; (2) prediction of chemical class membership using 43 binary neural net models; (3) unification of binary predictions to a final multiclass prediction using support vector machine (SVM). The dataset on which *Spec2Class* was trained, comprising of the in-house spectral library and publicly available data from the Mass Bank of North America.

Our predictor uses an ensemble of two machine learning algorithms. First, we performed binary classification using a convolutional neural network for each of the 43 classes. We then followed with a multiclass prediction using a support vector machine (SVM) model. This two-step prediction enabled us to combine all neural net outputs to one final class prediction (e.g., ‘this metabolite is a tritrepenoid’). Using ensemble classification enabled us to both identify class-specific features with a dedicated neural net, and consolidate all these neural nets to one unambiguous result. Apart from the final multiclass prediction, our method outputs a certainty parameter used to determine prediction confidence. Using the certainty parameter, one can filter low confidence results, attaining very high prediction accuracy. We made available a trained classifier that enables rapid prediction on uncharacterized mass spectra, runs on a stand-alone PC and works with both CPU and GPU architectures, though vastly accelerated if GPU is available. Implemented in Python, our open source method can be easily added to existing data processing pipelines, retrained with additional data, and customized to various research needs.

## Results

We devised a two-tiered classifier for plant secondary metabolite chemical class prediction, consisting of 43 neural-net binary classifiers, one for each class, followed by a final class determination with SVM (**Fig. 1b**). The neural networks consist of two 1D convolutional layers followed by three fully connected linear layers (**SI Fig. 1**). Each one of the binary neural-net binary classifiers accepts a binned spectrum as input and outputs an estimated probability of class membership. These 43 probabilities serve as input to the SVM model which provides the final class prediction. In addition, we introduced a certainty parameter for prediction quality control that allows the user to select only high confidence results (**Fig. 4, SI Fig. 3,4**)

### *Spec2Class* neural-net binary classifiers exhibit high classification accuracy

We sought to test the performance of both steps of the ensemble classifier. To test the first step which consists of neural net binary classifiers, we applied 15-fold cross validation to each of the 43 classifiers, setting a single set of hyperparameters for all, and varying only on the number of epochs for training (see Methods). Our model exhibited high accuracy in cross validation, with an average auROC of 0.91 across classes, ranging between 0.81 and 0.98 (**Fig. 2b**) on an average of 22 (6-38) training epochs. These results show that the binary neural net models were able to learn informative features in a short training process and that a single training setup could be applied to a wide range of chemical classes, even those with a small number of samples.

**Figure 2.**
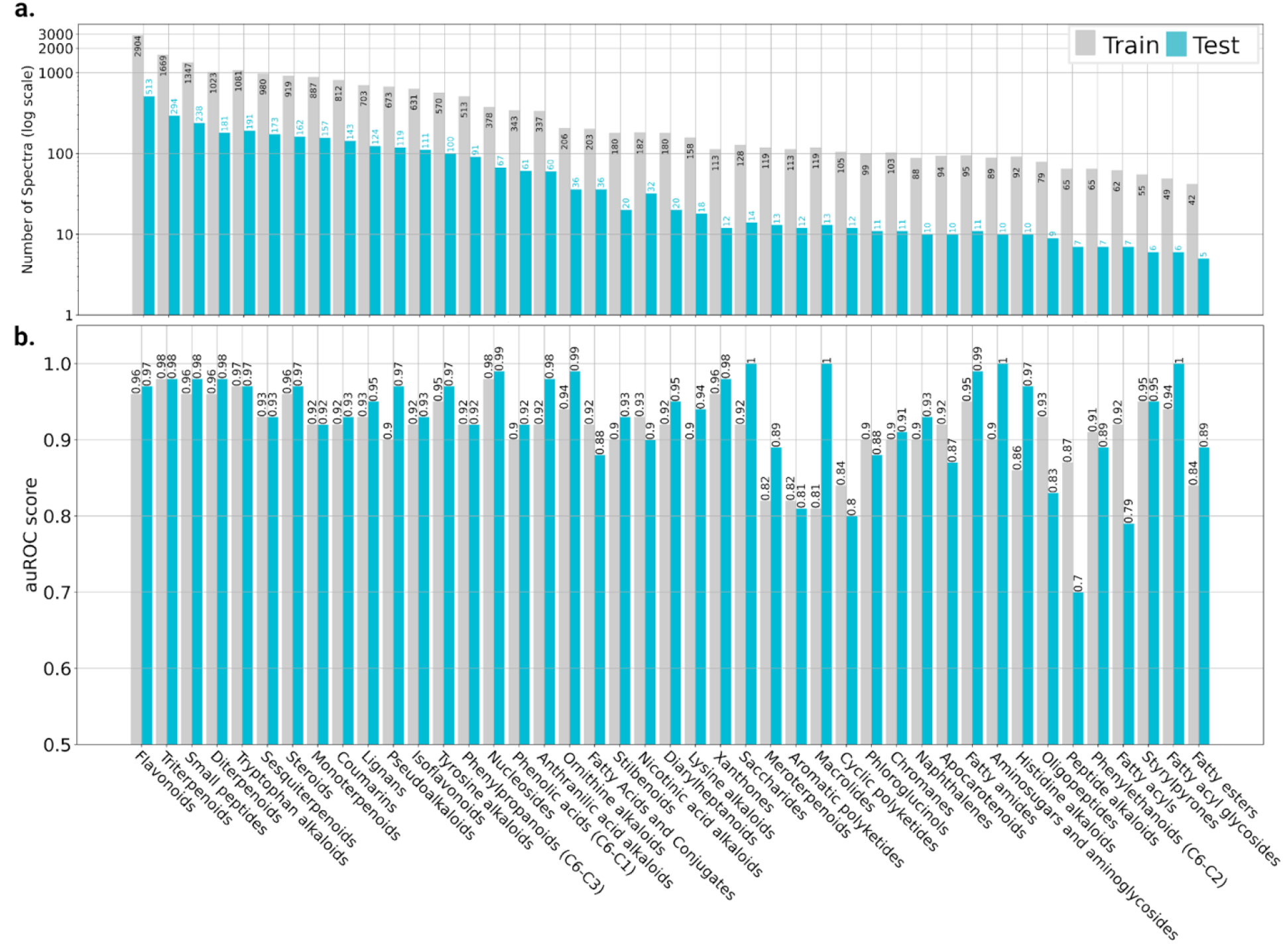
*Spec2Class* permits highly accurate binary classification. **(a)** Number of spectra from each class used for training and testing **(b)** Area under the receiver operating characteristic curve (auROC) scores for each one of the binary predictors. For each class we show both cross validation and test results.

Next, we sought to evaluate how our model generalizes by testing it on a held-out test set which had similar class and source (MoNA/in-house) distributions as the training set. We confirmed that this test set does not contain the same compounds as the training set using the unique InChiKey identifier. Our models showed similarly high accuracy on the test set, with an average auROC of 0.92. The best scoring superclass was *amino sugars and aminoglycosides* with an auROC of 0.99 and the worst was *peptide alkaloids* with 0.70 (**Fig. 2b**). Both classes had less than 100 spectra in the training set, and we therefore hypothesized that the difference in our ability to classify those stems from higher spectral variability in the *peptide alkaloids* group. To test this hypothesis, we calculated the averaged pairwise cosine similarity across mass spectra to determine intra-group similarity, whereby *amino sugars and aminoglycosides* received the highest value of 0.33 and *peptide alkaloids* received one of the lowest scores 0.04, supporting our hypothesis (**SI Table 4**). These results demonstrate that our binary models generalize well and may rely on a relatively small set of spectra for successful training.

To benchmark our binary models with an external tool, we compared to *SteroidXtract (12)*, which relies on a similar neural network, albeit on a smaller spectral range (50-200 Da). As *SteroidXtract* was trained to classify compounds to steroids or non-steroids, we compared it to our steroid classifier, using the same test set. In this comparison, *SteroidXtract* had an auROC score of 0.88, while the *Spec2Class* steroid binary predictor had an auROC of 0.97. Overall, our results show that training binary neural-net-based predictors on our data resulted in robust predictions for all classes, and in steroid classification that is superior to known predictors. We made our binary classification models for the 43 chemical classes publicly available in https://huggingface.co/VickiPol/binary_models.

### *Spec2Class* achieves accurate unambiguous classification using SVM

To receive a final concise multiclass prediction, avoiding ambiguous assignments to several classes, we used an SVM model with an radial basis function (RBF) kernel. Our SVM classifier receives as input, for each mass spectra sample, all 43 binary classification probabilities from the binary predictors (**Fig. 1b**). SVM re-estimates the class probabilities such that the class which best fits the spectra receives the highest score enabling us to greatly reduce ambiguity between binary predictions.

We evaluated the performance of the SVM model and selected hyperparameters using 20-fold cross validation. The best set of hyperparameters yielded an average training auROC score of 0.994. Upon adjusting the binary predictions with SVM, we compared the binary auROC classification (the SVM input) to the final classification (SVM output). The SVM auROC score was higher in 0.08 from the previous binary step (0.91). On the held-out test set, the average SVM auROC score was 0.94, an improvement over the binary classifier average (0.92). These results indicate that the SVM managed to capture additional information on the actual chemical class of each compound from the binary prediction vectors of all chemical classes, and thus improved the prediction.

We next tested these predictions on the held-out test, and achieved auROC scores ranging from 0.71 (phenylethanoids (C6-C2)) to 0.9988 (aminosugars and aminoglycosides), with the latter receiving the highest score both at this stage and the previous binary classification stage. These results showed that an additional SVM classification stage improves class determination, justifying the ensemble approach.

The ultimate test of our algorithm’s accuracy is its ability to correctly determine class membership. We therefore measured the percentage of instances where our top prediction was the correct one (top 1 accuracy score), or where the correct class assignment was within our top 3 predicted classes (top 3 accuracy score). On our training data, *Spec2Class* achieved a top 1 cross validation accuracy of 85.1% and a top 3 accuracy of 96.4%. On the held-out test set it achieved a top 1 accuracy of 73% and top 3 accuracy of 86% (**Fig. 3a**). These results show the robustness of our prediction and a good generalization of the SVM model.

**Figure 3:**
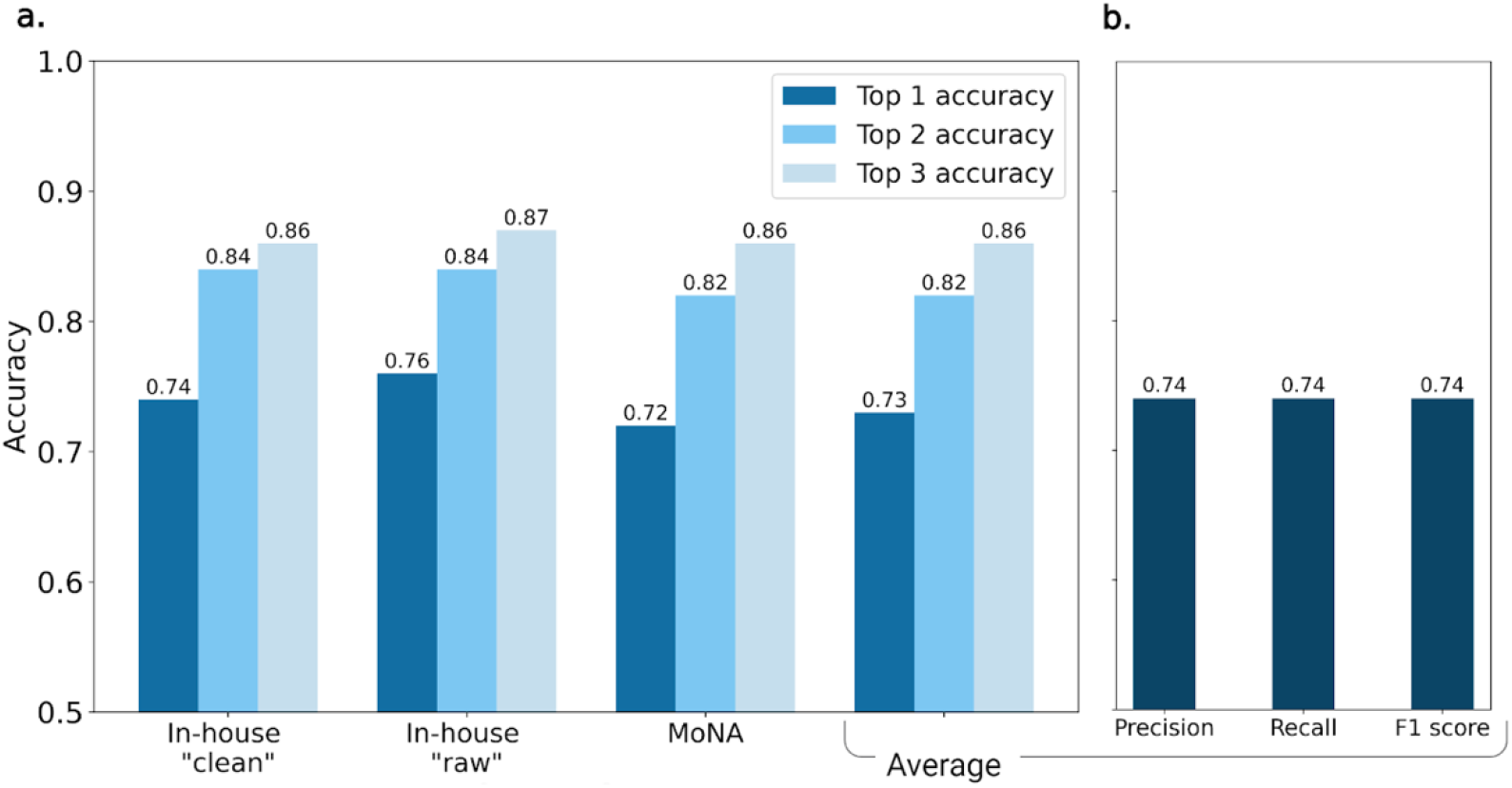
*Spec2Class* multiclass classification results. **(a)** Comparison of the accuracy results among the different data sources that compose the test set. For each data source the data preprocessing was performed differently, however these are not reflected in the results, demonstrating good generalization. **(b)** Average classification results of *Spec2Class* on the test set.

To examine possible differences in accuracy due to source or preprocessing, the prediction accuracy was checked separately for the three data sources present in the test set: “raw” spectra from in-house spectral library, “clean” spectra from in-house spectral library, and the spectra from MoNA. The highest held-out test set accuracy, 76% (**Fig. 3a**) originated from the noisiest source, the in-house raw spectra, the lowest (72%; **Fig. 3a**) from MoNA. The overall differences in the accuracy value between the sets is small, less than 4%, showing again the robustness of our prediction to data acquisition modes and post-acquisition processing. The average precision and recall values (**Fig. 3b)** are 74.5% and 73.6%, respectively, demonstrating good balance between quantity and quality of predictions.

### Prediction confidence parameter (p1-p2)

To quantify the confidence of *Spec2Class* predictions, we sought to provide a metric that may be used to filter out low confidence predictions (**SI Fig. 3,4; SI Table 5**). To this end, we analyzed the output probability vectors given for each sample by the SVM model and observed that the difference between the prediction probability of the top class (p1) and the prediction probability of the second-best class (p2) was highly correlated with the top 1 prediction accuracy. We termed this prediction margin p1-p2 and quantified the test set accuracy achieved at every selected threshold. For example, samples with p1-p2 value higher than 0.9, were predicted with 93% accuracy on the held-out test set (**SI Fig. 3**).

Higher confidence prediction comes with a cost of the reduction in the number predictions. The p1-p2 value is given for each output, enabling the user to decide on the tradeoff between the number of predicted samples and the level of prediction confidence, similarly to false discovery rate (SI Fig. 4).

### *Spec2Class* shows high accuracy on independent data sets

To comprehensively assess *Spec2Class’s* performance, a thorough benchmarking was conducted against *CANOPUS* (11) utilizing an external validation set. This set was derived from three independent sources acquired through LC-ESI-QToF positive mode. Firstly, 175 spectra from a publicly available dataset from Sam Sik Kang (14) was selected to mitigate biases associated with in-house data processing. Secondly, 102 spectra from the in-house ‘superpool’ set (15), originating from a complex biological matrix spiked with standards (Supplementary Information), was chosen due to the common ‘matrix effect’ characterized by higher noise levels in the spectra arising from the mixture. Lastly, 55 spectra from the ‘Tomato BILs’ (tomato backcross inbred lines) set (16), an in-house compilation of manually labeled and curated spectra from two developmental stages of tomato fruit, was included. In contrast to the ‘superpool’ set, no standard spiking was performed during sample preparation. ‘Tomato BILs’ set was chosen to further challenge the model and assess its performance on data obtained from a biological experiment.

To facilitate a fair evaluation using the p1-p2 threshold, we restricted our comparison to identical spectra, even though confidence filtering is not available in *CANOPUS* (**SI Table 6)**. The reduced number of spectra predicted by *CANOPUS* is attributed to prolonged computation times required for heavier compounds, prompting the need for time constraints to obtain at least partial results. In the absence of p1-p2 confidence filter, *CANOPUS* exhibits a slight superiority over *Spec2Class* by 4% in precision (0.75 compared to 0.79). However, *Spec2Class* surpasses *CANOPUS* by 3% in recall and lags by only 1% in F1 score. Upon applying a p1-p2 threshold higher than 0.5, *Spec2Class* outperforms *CANOPUS*.

Beyond demonstrating comparable results to state-of-the-art model, we reaffirm the fidelity of the p1-p2 confidence parameter on additional external sets (**Fig. 4c** and **SI Tables 5 and 6**). These outcomes underscore the high classification scores of *Spec2Class* in the context of plant-derived compounds, despite its independence from molecular fingerprinting and chemical formulae. The option to filter low-confidence predictions reduces the number of predictions but also significantly enhances both precision and recall.

**Fig 4.**
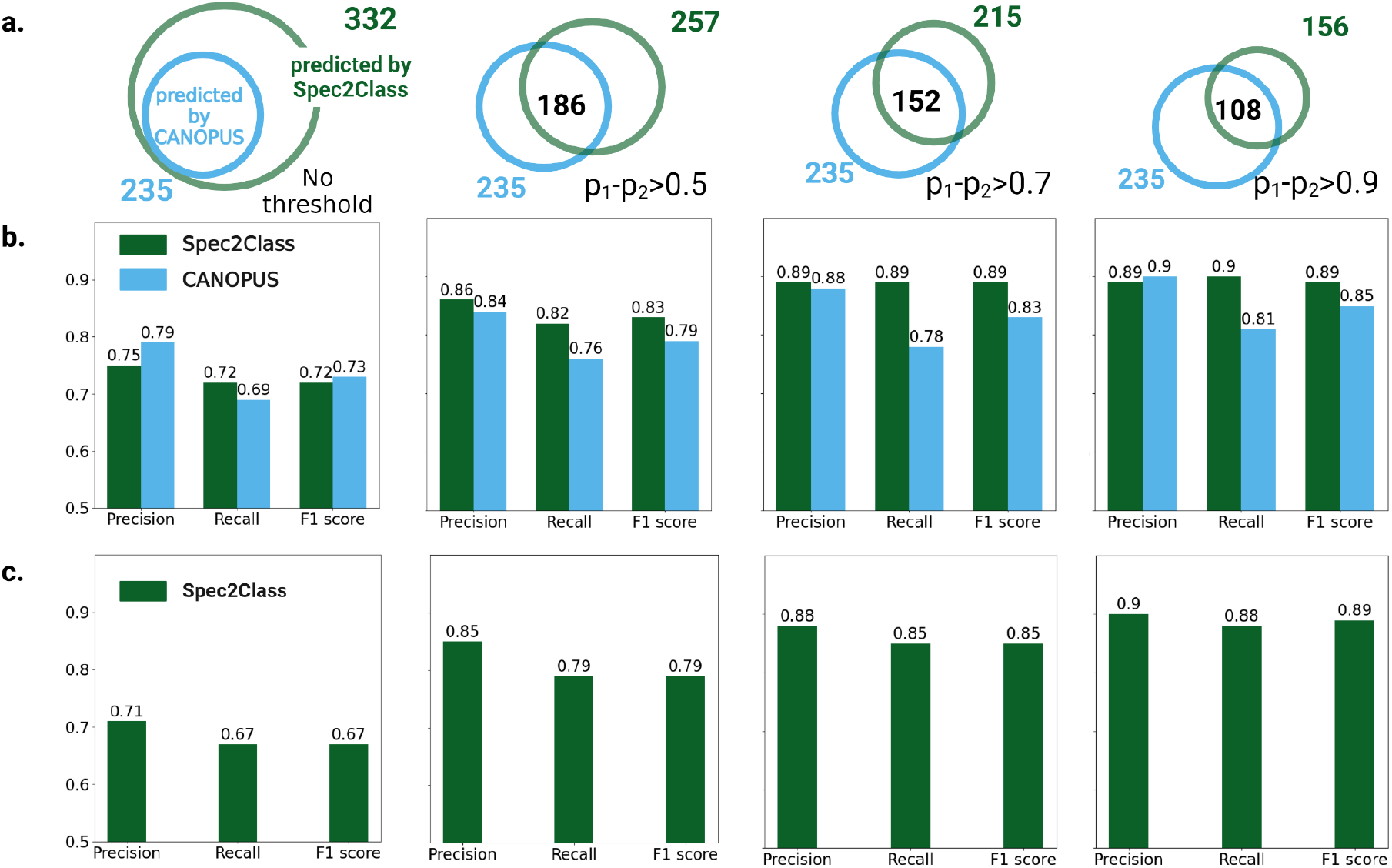
Benchmarking with *CANOPUS* (11) on external validation set. **(a)** Graphical representation of the overall number of predictions obtained by each model and their intersection. *Spec2Class* prediction confidence can be increased with result filtration based on p1-p2 threshold. **(b)** Comparison of *Spec2Class* results to *CANOPUS* on both predicted spectra (intersection). *Spec2Class* outperforms *CANOPUS* when applying p1-p2 threshold higher than 0.5. **(c)** *Spec2Class* results on the entire external validation set with the application of p1-p2 threshold.

## Discussion

*Spec2Class* is an ensemble classifier, which predicts chemical superclass directly from MS/MS spectra. It was trained in a supervised manner on a rich and diverse standard spectra library with almost 8,000 chemical standards of plant secondary metabolites and ∼14,000 publicly available spectra. *Spec2Class* resulted with a top 1 accuracy and F1 score of 73% on the test set. Its performance was almost similar between the different data sources, processing methods and data acquisition modes. Surprisingly, the accuracy was higher for unprocessed, ‘raw’ spectra of chemical standards compared to the ‘clean’ ones, which passed an extensive process of noise reduction. This result proves the robustness of the model and its generalizability and suggests that some information which is valuable for prediction may be lost in the cleaning process.

There was a clear class imbalance in our data set, whereby for all classes the number of negative samples surpassed the number of positives. To overcome this challenge and reduce overfitting, we applied several regularization techniques, including augmentation by spectral subsampling, neural net dropout and loss correction (Supplementary Information). The regularization allowed us to improve the training process and to use a similar training framework for all the binary predictors, regardless of the number and diversity of the positive examples. It demonstrates that the binary model architecture and training framework are a good fit for this prediction task and can be easily used for additional chemical classes, with additional data. The script, implemented in Python, is highly modular and openly available for future community development and model expansion.

Compared to *CANOPUS (11), Spec2Class* classification scores are higher with the use of the confidence parameter and comparable without. All the data sets contained plant sourced compounds, highlighting the capacity of *Spec2Class* in predicting plant secondary metabolites classes. Apart from differences in their predictive performance, *Spec2Class* has several advantages: (a) It can run locally, without a webserver; (b) The speed of prediction is indifferent with respect to the precursor mass and can be accelerated with the use of GPU; (c) It outputs with the prediction a confidence parameter that estimates its chance to be correct; (d) It is implemented in Python and well documented, and is thus highly modular and can be modified, extended and incorporated to data processing pipelines.

*Spec2Class* was trained on positive ionization mode spectra obtained by an LC-ESI-QToF setup, a common metabolomics technology platform. A different setup will likely result with different fragmentation patterns for similar compounds, thus the prediction quality using the same training set is expected to be lower for other types of instruments. The difference in fragmentation patterns also applies to positive versus negative ionization modes; thus the usage of *Spec2Class* is currently limited to the positive ionization mode. Notwithstanding, *Spec2Class* could be easily re-trained in a similar manner on a data set of spectra obtained by negative ionization mode or other instrumentation, with likely similar accuracy. Positive mode remains the most common ionization mode in publicly available data, however in non-targeted experiments, samples are often obtained by both charging modes. The different fragmentations that are produced can provide additional and complementary information that helps in compound identification. We believe that the usage of two models, one trained for positive and one for negative ionization mode, could improve the confidence of prediction.

Resulting with an accurate prediction for 43 major chemical super classes in plant metabolites, *Spec2Class* is a reliable and useful tool for plant metabolomics. We anticipate that *Spec2Class* may also prove beneficial for metabolites originating from other organisms, simply because not all the chemical classes are unique to plants (for example steroids, small peptides, saccharides, etc.). While half of the data set used to train *Spec2Class* consists of spectra from MoNA and lacks information about the organism, it is likely that the training set comprises spectra from various organisms. *Spec2Class* does not solve the task of accurate metabolite structure elucidation. However, the knowledge of the most probable compound superclass could facilitate this task. In addition, *Spec2Class* can be highly useful for studies that do not require full annotation but can rely on the comparison of chemical family abundance across samples. Moreover, it could also serve as the initial phase of metabolomics experiments examination providing a ‘bird’s eye’ view of metabolic changes at the class level.

As an automatic tool, the model will provide a prediction of one of the 43 trained chemical classes, for any input. The 43 chemical classes cover a big portion of the plant chemical space, but not all of it. Thus, the prediction might be misleading for compounds that do not belong to one of these classes. A key feature in the model is the prediction certainty parameter (p1-p2). The p1-p2 parameter allows the user to consider only the most confident results, thus reducing false discovery risk. We assume that compounds that do not belong to one of the 43 predicted chemical classes will receive a low p1-p2 value and thus be filtered out when employing this filter. *Spec2Class* is an important first step towards a future expansion of this prediction setup to additional chemical classes, instrument setups and ionization modes. It demonstrates the ability of deep learning models to assist in mass spectrometry experiments, and we expect the accuracy of these models to vastly improve in accuracy and scope with additional data points.

## Materials and Methods

### 1. Input data

The data used in this work is composed of positive ionization mode spectra sourced form (a) in-house spectral library, obtained through data independent acquisition (DIA) (15); (b) the Mass-bank of North America (MoNA; https://mona.fiehnlab.ucdavis.edu/), obtained through data dependent acquisition (DDA). We split the data into training and test sets to assure proper benchmarking and scoring of our method, consisting of 23,769 and 3,847 spectra, respectively. To ensure similar distributions of chemical super classes and data sources in each set, we assigned a test set with 15% and 10% of samples from each class for super classes with over 200 and fewer than 200 spectra, respectively (**Fig. 2a**)

#### 1.1. In-house spectra data acquisition

LC-MS data of chemical standards pools were experimentally acquired and computationally processed as in *Weizmass* (15), with the exception of the additional spectra cleaning step detailed below. For each compound in the in-house library, we used two types of spectra in our data set; one, labeled as “clean” and second labeled as “raw”. Both types of spectra went through the standard procedure of mass spectrometry data analysis resulting in a peak table. Unlike “raw” spectra, the “clean” spectra were filtered by retaining the top 30 most intense MS2 ions, with preference to ions for which the chemical formula could be predicted using the corresponding isotopic pattern. We chose to use both versions of spectra for better algorithm generalization. The “clean” spectra have a lower number of fragments compared to the “raw” version. “Clean” spectra might miss fragments that have been wrongly filtered, while the “raw” version might contain excess “noise” fragments. Our training algorithm promised that both types of spectra from similar compounds will appear either in the training set or in the test set, but not in both. This rule was applied to the cross-validation partitioning of the training set, promising that in each fold spectra from a similar compound will appear either in training or in validation set.

### 2. Spectra labeling

We labeled the training and test sets with *NPClassifier* (13) based on the SMILES string of each compound. We demanded that chemical classes in our dataset have at least 40 spectra, to ensure proper training and testing, resulting in a total of 43 chemical classes. *NPClassifier* was chosen for the task of spectra labeling due to its higher specificity for natural product ontology.

### 3. Binary prediction

We trained 43 binary classifiers, one for each superclass, followed by a single SVM classifier for consolidating the binary results. We binned the input spectra into m/z bins and vectorized an input list of m/z and intensity pairs into a binned vector of length 5000, representing masses between 50 to 550 Da with a resolution of 0.1 Da, where every value in the vector corresponds to the intensity of that fragment. For m/z values that fall into the same bin, we assigned the higher intensity value. The lower mass limit of 50 Da was chosen according to the mass detector specifications. The upper mass limit was set at 550 Da to assure proper representation of all mass features and includes more than 90% of the fragments in the data set (**SI. Table 1**). We normalized these binned spectra as previously described (Supplementary Information; (12)).

#### 3.1. Neural net architecture

We trained 43 binary classifiers, one for every chemical class. Each such classifier is a neural net model with an architecture based on *SteroidXtract (12)* consisting of two one-dimensional convolutional layers followed by three fully connected linear layers (**SI Fig. 2**). We redesigned our convolutional layers to fit the size of the input vector. Each of the 43 binary models, one per chemical superclass, outputs a predicted probability that this spectrum belongs to that class.

#### 3.2. Cross validation

We split the training set into 15 folds, ensuring that the frequency of positive samples (those belonging to a certain class, per binary prediction model) was similar in the training and validation sets (**SI Table 2**). Since some chemicals were profiled more than once, often with different spectra, we made sure that the same chemical compounds did not appear in both the training and the test sets. To this end, we used the GroupKFold function from the scikit-learn Python package (17). We grouped compounds using their ‘Chemical_ID’ parameter, which is the first 1/3 of the international chemical identifier key (InChIKey) and represents only the chemical structure, without spatial information. This allowed us to differentiate spectra from different molecules, while ignoring differences between spectra produced by stereoisomers. We assessed the binary model’s performance using the area under the receiver operating characteristic curve (auROC score) and used the maximal auROC out of all training epochs, averaged across all folds to determine the optimal number of training epochs for each class, as well as the optimal hyperparameters. Our final model used the following values: lr = 0.0001, batch_size = 128, dropout_conv = 0.95, dropout_linear = 0.95.

### 4. Multiclass prediction

The second stage of our predictor used the outputs of all binary classifiers to unambiguously classify spectra to a single chemical super class. We trained a support vector machine (SVM), provided by scikit-learn package (17) with a radial basis function (RBF) kernel, a gamma coefficient of 0.01 and C equal to 100. The choice of the kernel function, gamma and C coefficients were based on the results of a hyperparameter search on 20-fold cross validation, selecting the hyperparameters that achieved the highest average accuracy. Accuracy was calculated as the number of correct classifications divided by the total number of samples. The input for the SVM model is a matrix of size (samples X super classes). Each matrix row represents one sample. Column values are the results of all superclass binary models on that sample. The output of the SVM is a vector of class labels, with the final prediction of the chemical superclass. To estimate top 2, 3 accuracy values on the test set results, we used the ‘predict_proba’ method from scikit-learn package (17) to receive an output probability vector for each sample. These vectors were also used to calculate the auROC scores for each class.

### 5. Comparison to other models

We compared *Spec2Class* to *SteroidXtrac*t (12) using the auROC score, applied to the *Spec2Class* classification of steroids. We compared *Spec2Class* to *CANOPUS (11)* using its *SIRIUS GUI* version 5.8.1 (see full command in Supplementary Information). *CANOPUS* prediction was preceded by the *SIRIUS (18)* module for molecular fingerprint prediction and by the *CSI:FingerID* (7) module for molecular fingerprint prediction. As *Spec2Class* requires no molecular formula, nor structural information for its prediction, we compared on equal terms and removed the formula and molecular string representations from the input. The compounds tested in this comparison were not a part of the *Spec2Class* training set spectra.

Some of the spectra with precursor masses higher than 850 Da were excluded from the comparison due to extremely long computation time of fragmentation trees that are a part of the *CANOPUS* algorithm. Using *SIRIUS* GUI, we set a ‘Tree timeout’ and ‘Compound timeout’ to 10 minutes to allow the calculation to complete without a prior exclusion of compounds heavier than 850 Da. Both algorithms were tested on three independent data sets of spectra from a plant source, obtained by positive ionization mode and LC-ESI-QToF setup: (a) Sam Sik Kang Legacy library CCMSLIB00010007697 to CCMSLIB00010008032 in the spectral library of GNPS (19); (b) the ‘spiked superpool’ set (15), (c) Tomato BILs set (16).

## Notes

### Competing Interest Statement

The authors have declared no competing interest.

