## Supplementary Information for "Spec2Class: Accurate Prediction of Plant Secondary Metabolite Class using Deep Learning"

#### Spectrum intensity normalization

For each spectrum, the intensities were normalized as follows:

$$x_{i_{normalized}} = \sqrt{\frac{100 * x_i}{max\ (x)}}$$

$$x_{i_{normalized}} - normalized\ intensity\ value\ of\ a\ single\ fragment$$

$$x_{i_{normalized}} - intensity\ value\ of\ a\ single\ fragment,$$

$$(x)\ - maximal\ intensity\ value\ within\ a\ spectrum$$

#### Improving the learning process of the binary classifiers

To tackle the common problem of the imbalanced set as well as the overfitting to the training data several actions were taken:

##### Data augmentation

Augmentation is a regularization technique used for model generalization. In practice, this is the process of data set supplementation with ‘artificial’ samples from the existing data set. Good augmentation mimics real life samples that might contain flaws, forcing the model to extract informative and generalized features rather than noise or distortions in the training process. To apply this strategy in mass spectrometry, the main idea was to mimic the common absence of low intensity fragments in a spectrum. To this end, for each spectrum in the data set, we generated a series of ‘artificial’ spectra by subsampling a set portion of fragments in the spectrum with a weighted probability proportional to the relative intensity of the fragment. Each such sampling process was repeated a number of times. For example: if a spectrum contained 20 fragments and was sampled at 80%, the generated spectra will have 16 fragments and the fragments with higher relative intensity values will be those with the higher probability to be sampled. Each spectrum was subsampled 9 times: 4 at 80%, 3 at 70% and 2 at 60%. In order to compensate for the dataset class imbalance, the amount of added augmented spectra was limited to the number (fill the exact number) which is equal to 1.5 multiplied by the number of spectra in the largest class. This way the portion of added spectra was bigger for smaller classes compared to the larger classes.

**Weighted loss function**

In the binary model we used the binary cross entropy loss function as was used in *SteroidXtract (1)*. To improve the training process, and address the class imbalance, we applied the weighted BCEloss function from PyTorch python library (2). The weights were calculated for each sample type (positive or negative, e.g., is or isn't a triterpenoid), according to its portion in each batch in the training loop. The weight for each type of sample was equal to 1 / sample portion. The loss value of each sample was multiplied by its weight, such that samples with higher portions in the training set received a lower per-sample loss value. The use of weights helped to prevent the bias towards the type of samples that were most frequent, thus improving the learning process.

**Increasing the dropout portion**

The term "dropout" refers to dropping out the nodes (input and hidden layer) in a neural network. All the forward and backwards connections with a dropped node are temporarily removed, thus creating a new network architecture from the parent network, with a reduced number of parameters. The nodes are dropped by a dropout probability value that is set in advance. This is another common method of regularization that was especially useful in our case.

The neural net architecture contains two dropout layers in the training phase: (a) after the second convolutional layer; (b) after the second linear layer (FIGURE REFERENCE). Based on the hyperparameter search, both dropout probabilities were set to 0.95.

##### Correlations

We checked possible correlations between number of samples in class, final loss values, best auROC scores and number of training epochs (Supplementary Fig 3). The Spearman correlation was calculated based on average cross validation results of all the binary classifiers using scikit-learn package (3).

Class size had resulted in the strongest correlation with the optimal epoch number ($R=0.72$, $p<0.05$; Supplementary Fig. 3). auROC score and loss value were the only pair of parameters for which the correlation was not significant ($p>0.05$).

##### CANOPUS command

```
config --IsotopeSettings.filter=true --FormulaSearchDB= --Timeout.secondsPerTree=600
--FormulaSettings.enforced=HCNOP --Timeout.secondsPerInstance=600
--AdductSettings.detectable=[[M-H4O2+H]+,[M+Na]+,[M+H3N+H]+,[M+H]+,[M+K]+,[M-H2O+H]+
] --UseHeuristic.mzToUseHeuristicOnly=650 --AlgorithmProfile=qtof
--IsotopeMs2Settings=IGNORE --MS2MassDeviation.allowedMassDeviation=10.0ppm
--NumberOfCandidatesPerIon=1 --UseHeuristic.mzToUseHeuristic=300
--FormulaSettings.detectable=B,Cl,Br,Se,S --NumberOfCandidates=10
--AdductSettings.enforced=, --AdductSettings.fallback=[[M+Na]+,[M+H]+,[M]+,[M+K]+]
--FormulaResultThreshold=true --RecomputeResults=false formula fingerprint canopus
```

#### Superpool data set preparation

###### Complex biological matrix preparation

An equal (1:1) extract mixture of *Arabidopsis thaliana* leaves and tomato (*Solanum lycopersicum*; cv. M82) fruit red skin (referred as 'superpool') was used to prepare the sample pools. Extract preparation: frozen leaves of *A. thaliana* from 3.5-week-old wild-type Columbia-0 plants were ground and extracted with an extraction solution (80% methanol and 0.1% formic acid) at a ratio of 1:3 (w/v). Samples were sonicated for 20 min in a bath sonicator, vortexed and centrifuged for 12 min at 14,000g, and the supernatant was filtered through 0.22 mm polyvinylidene difluoride filters. Red skin of tomato fruit was extracted using 100% methanol (1:3 (w/v)).

**Spiking chemical standards into the ‘superpool’**

Randomly picked metabolite standards were injected into the ‘superpool’ samples. The final concentration of the chemical standards was about 7 mg /ml for each. The data processing was performed according to the *WeizMass* workflow described by N. Shachaf et al. (4).

### Supplementary Tables

Supplementary Table 1 : Distribution of mass to charge ratio (m/z) values in training data set

| Total Number of Spectra in Training Set | Maximal Fragment m/z [Da] | Mean Fragment m/z [Da] | 5th Percentile of Fragment m/z [Da] | 10th Percentile of Fragment m/z [Da] | 50th Percentile of Fragment m/z [Da] | 90th Percentile of Fragment m/z [Da] | 95th Percentile of Fragment m/z [Da] |
|---|---|---|---|---|---|---|---|
| 23769 | 2994 | 112 | 67 | 93 | 225 | 507 | 623 |

Supplementary Table 2: Cross validation average results of the binary classifiers

| Class | auROC score | TP | TN | FN | FP | Number of positive samples in a fold | Number of negative samples in a fold | Best number of epochs | loss |
|---|---|---|---|---|---|---|---|---|---|
| **Aminosugars and aminoglycosides** | 0.899 | 73 | 2303 | 12 | 49 | 85 | 2352 | 11 | 0.08 |
| **Anthranilic acid alkaloids** | 0.92 | 157 | 1429 | 69 | 113 | 227 | 1542 | 22 | 0.31 |
| **Apocarotenoids** | 0.918 | 64 | 1526 | 25 | 39 | 88 | 1565 | 20 | 0.15 |
| **Aromatic polyketides** | 0.823 | 51 | 1435 | 54 | 119 | 105 | 1554 | 17 | 0.28 |
| **Chromanes** | 0.9 | 54 | 1496 | 30 | 68 | 83 | 1564 | 20 | 0.2 |
| **Coumarins** | 0.921 | 282 | 1373 | 83 | 126 | 365 | 1498 | 39 | 0.37 |
| **Cyclic polyketides** | 0.835 | 39 | 1464 | 41 | 101 | 80 | 1565 | 15 | 0.24 |
| **Diarylheptanoids** | 0.916 | 119 | 1493 | 48 | 64 | 167 | 1557 | 29 | 0.25 |
| **Diterpenoids** | 0.957 | 337 | 1401 | 59 | 68 | 396 | 1469 | 36 | 0.27 |
| **Fatty Acidsand Conjugates** | 0.925 | 75 | 1491 | 52 | 66 | 127 | 1557 | 20 | 0.19 |
| **Fatty acyls** | 0.922 | 24 | 1521 | 18 | 49 | 43 | 1570 | 18 | 0.17 |
| **Fatty acyl glycosides** | 0.943 | 28 | 1550 | 9 | 21 | 38 | 1571 | 15 | 0.08 |
| **Fatty amides** | 0.953 | 50 | 1506 | 19 | 61 | 68 | 1568 | 11 | 0.12 |
| **Fatty esters** | 0.842 | 8 | 1559 | 16 | 12 | 24 | 1572 | 17 | 0.1 |
| **Flavonoids** | 0.961 | 528 | 1151 | 61 | 120 | 589 | 1271 | 45 | 0.28 |
| **Histidine alkaloids** | 0.864 | 31 | 1367 | 26 | 201 | 57 | 1568 | 6 | 0.3 |
| **Isoflavonoids** | 0.917 | 269 | 1370 | 76 | 144 | 345 | 1514 | 30 | 0.4 |
| **Lignans** | 0.933 | 267 | 1436 | 89 | 71 | 356 | 1508 | 32 | 0.3 |
| **Lysine alkaloids** | 0.903 | 75 | 1436 | 36 | 123 | 111 | 1559 | 17 | 0.23 |
| **Macrolides** | 0.813 | 39 | 1469 | 36 | 97 | 75 | 1566 | 16 | 0.24 |
| **Meroterpenoids** | 0.82 | 51 | 1480 | 53 | 82 | 104 | 1562 | 34 | 0.27 |
| **Monoterpenoids** | 0.92 | 278 | 1386 | 81 | 121 | 359 | 1507 | 32 | 0.32 |
| **Naphthalenes** | 0.895 | 39 | 1502 | 34 | 64 | 73 | 1566 | 15 | 0.18 |
| **Nicotinic acid alkaloids** | 0.932 | 79 | 1466 | 39 | 93 | 118 | 1558 | 19 | 0.21 |
| **Nucleosides** | 0.982 | 191 | 1491 | 27 | 58 | 219 | 1549 | 17 | 0.16 |
| **Oligopeptides** | 0.931 | 24 | 1543 | 30 | 26 | 54 | 1569 | 12 | 0.11 |
| **Ornithine alkaloids** | 0.94 | 101 | 1479 | 37 | 77 | 138 | 1556 | 19 | 0.19 |
| **Peptide alkaloids** | 0.869 | 17 | 1508 | 21 | 62 | 37 | 1570 | 11 | 0.19 |
| **Phenolic acids (C6-C1)** | 0.9 | 14 | 1502 | 14 | 45 | 28 | 1547 | 18 | 0.12 |
| **Phenylethanoids (C6-C2)** | 0.912 | 2 | 1562 | 8 | 3 | 9 | 1565 | 23 | 0.03 |
| **Phenylpropanoids (C6-C3)** | 0.915 | 34 | 1403 | 11 | 126 | 46 | 1529 | 26 | 0.22 |
| **Phloroglucinols** | 0.898 | 64 | 1470 | 29 | 93 | 93 | 1563 | 17 | 0.2 |
| **Polycyclic aromatic polyketides** | 0.84 | 43 | 1474 | 48 | 90 | 91 | 1564 | 18 | 0.33 |
| **Pseudoalkaloids** | 0.898 | 211 | 1402 | 130 | 121 | 341 | 1524 | 36 | 0.46 |
| **Saccharides** | 0.916 | 56 | 1519 | 15 | 47 | 72 | 1566 | 12 | 0.23 |
| **Sesquiterpenoids** | 0.935 | 301 | 1412 | 86 | 64 | 387 | 1477 | 28 | 0.27 |
| **Small peptides** | 0.964 | 328 | 1376 | 58 | 102 | 387 | 1478 | 29 | 0.26 |
| **Steroids** | 0.959 | 295 | 1439 | 61 | 68 | 356 | 1507 | 28 | 0.21 |
| **Stilbenoids** | 0.903 | 115 | 1458 | 51 | 99 | 166 | 1557 | 23 | 0.27 |
| **Styrylpyrones** | 0.953 | 39 | 1495 | 13 | 74 | 52 | 1569 | 21 | 0.14 |
| **Triterpenoids** | 0.976 | 365 | 1395 | 49 | 54 | 413 | 1449 | 21 | 0.17 |
| **Tryptophan alkaloids** | 0.971 | 337 | 1391 | 45 | 92 | 382 | 1483 | 36 | 0.22 |
| **Tyrosine alkaloids** | 0.947 | 265 | 1420 | 70 | 109 | 335 | 1529 | 32 | 0.29 |
| **Xanthones** | 0.956 | 90 | 1494 | 25 | 69 | 115 | 1562 | 23 | 0.16 |

Supplementary Table 3: Test average results of the binary classifiers

| Class | auROC score | TP | TN | FN | FP | Number of positive samples | Number of negative samples in a |
|---|---|---|---|---|---|---|---|
| **Aminosugars and aminoglycosides** | 0.999 | 10 | 3794 | 0 | 39 | 10 | 3833 |
| **Anthranilic acid alkaloids** | 0.985 | 70 | 3549 | 6 | 218 | 76 | 3767 |
| **Apocarotenoids** | 0.872 | 6 | 3796 | 8 | 33 | 14 | 3829 |
| **Aromatic polyketides** | 0.808 | 11 | 3660 | 22 | 150 | 33 | 3810 |
| **Chromanes** | 0.912 | 10 | 3732 | 3 | 98 | 13 | 3830 |
| **Coumarins** | 0.935 | 157 | 3375 | 40 | 271 | 197 | 3646 |
| **Cyclic polyketides** | 0.801 | 6 | 3637 | 9 | 191 | 15 | 3828 |
| **Diarylheptanoids** | 0.949 | 19 | 3686 | 7 | 131 | 26 | 3817 |
| **Diterpenoids** | 0.975 | 248 | 3418 | 32 | 145 | 280 | 3563 |
| **Fatty Acids and Conjugates** | 0.879 | 23 | 3724 | 18 | 78 | 41 | 3802 |
| **Fatty acyl glycosides** | 0.998 | 6 | 3821 | 1 | 15 | 7 | 3836 |
| **Fatty acyls** | 0.791 | 4 | 3751 | 5 | 83 | 9 | 3834 |
| **Fatty amides** | 0.988 | 10 | 3786 | 2 | 45 | 12 | 3831 |
| **Fatty esters** | 0.886 | 1 | 3814 | 4 | 24 | 5 | 3838 |
| **Flavonoids** | 0.972 | 752 | 2788 | 57 | 246 | 809 | 3034 |
| **Histidine alkaloids** | 0.974 | 7 | 3660 | 3 | 173 | 10 | 3833 |
| **Isoflavonoids** | 0.931 | 118 | 3375 | 35 | 315 | 153 | 3690 |
| **Lignans** | 0.949 | 130 | 3567 | 26 | 120 | 156 | 3687 |
| **Lysine alkaloids** | 0.939 | 17 | 3666 | 7 | 153 | 24 | 3819 |
| **Macrolides** | 0.996 | 13 | 3799 | 1 | 30 | 14 | 3829 |
| **Meroterpenoids** | 0.89 | 10 | 3728 | 5 | 100 | 15 | 3828 |
| **Monoterpenoids** | 0.915 | 138 | 3401 | 35 | 269 | 173 | 3670 |
| **Naphthalenes** | 0.93 | 8 | 3714 | 5 | 116 | 13 | 3830 |
| **Nicotinic acid alkaloids** | 0.9 | 22 | 3659 | 10 | 152 | 32 | 3811 |
| **Nucleosides** | 0.988 | 62 | 3648 | 6 | 127 | 68 | 3775 |
| **Oligopeptides** | 0.828 | 3 | 3797 | 7 | 36 | 10 | 3833 |
| **Ornithine alkaloids** | 0.99 | 35 | 3687 | 1 | 120 | 36 | 3807 |
| **Peptide alkaloids** | 0.702 | 4 | 3759 | 4 | 76 | 8 | 3835 |
| **Phenolic acids (C6-C1)** | 0.918 | 43 | 3641 | 19 | 140 | 62 | 3781 |
| **Phenylethanoids (C6-C2)** | 0.885 | 2 | 3816 | 18 | 7 | 20 | 3823 |
| **Phenylpropanoids (C6-C3)** | 0.923 | 93 | 3421 | 30 | 299 | 123 | 3720 |
| **Phloroglucinols** | 0.877 | 10 | 3689 | 4 | 140 | 14 | 3829 |
| **Polycyclic aromatic polyketides** | 0.741 | 7 | 3716 | 13 | 107 | 20 | 3823 |
| **Pseudoalkaloids** | 0.973 | 118 | 3523 | 11 | 191 | 129 | 3714 |
| **Saccharides** | 0.996 | 11 | 3798 | 3 | 31 | 14 | 3829 |
| **Sesquiterpenoids** | 0.935 | 188 | 3493 | 45 | 117 | 233 | 3610 |
| **Small peptides** | 0.979 | 228 | 3374 | 11 | 230 | 239 | 3604 |
| **Steroids** | 0.974 | 151 | 3534 | 23 | 135 | 174 | 3669 |
| **Stilbenoids** | 0.934 | 23 | 3674 | 5 | 141 | 28 | 3815 |
| **Styrylpyrones** | 0.951 | 7 | 3715 | 2 | 119 | 9 | 3834 |
| **Triterpenoids** | 0.981 | 298 | 3411 | 39 | 95 | 337 | 3506 |
| **Tryptophan alkaloids** | 0.972 | 201 | 3437 | 24 | 181 | 225 | 3618 |
| **Tyrosine alkaloids** | 0.975 | 99 | 3526 | 12 | 206 | 111 | 3732 |
| **Xanthones** | 0.984 | 15 | 3720 | 4 | 104 | 19 | 3824 |

Supplementary Table 4: Cosine similarity of spectra within the different chemical classes

| Class | Cosine similarity within class |
|---|---|
| **All vs all** | 0.05 |
| **Aminosugars and aminoglycosides** | 0.33 |
| **Anthranilic acid alkaloids** | 0.04 |
| **Apocarotenoids** | 0.25 |
| **Aromatic polyketides** | 0.05 |
| **Chromanes** | 0.02 |
| **Coumarins** | 0.08 |
| **Cyclic polyketides** | 0.06 |
| **Diarylheptanoids** | 0.11 |
| **Diterpenoids** | 0.14 |
| **Fatty Acids and Conjugates** | 0.08 |
| **Fatty acyl glycosides** | 0.23 |
| **Fatty acyls** | 0.15 |
| **Fatty amides** | 0.17 |
| **Fatty esters** | 0.15 |
| **Flavonoids** | 0.05 |
| **Histidine alkaloids** | 0.10 |
| **Isoflavonoids** | 0.03 |
| **Lignans** | 0.08 |
| **Lysine alkaloids** | 0.08 |
| **Macrolides** | 0.02 |
| **Meroterpenoids** | 0.10 |
| **Monoterpenoids** | 0.14 |
| **Naphthalenes** | 0.03 |
| **Nicotinic acid alkaloids** | 0.07 |
| **Nucleosides** | 0.09 |
| **Oligopeptides** | 0.01 |
| **Ornithine alkaloids** | 0.11 |
| **Peptide alkaloids** | 0.04 |
| **Phenolic acids (C6-C1)** | 0.09 |
| **Phenylethanoids (C6-C2)** | 0.11 |
| **Phenylpropanoids (C6-C3)** | 0.15 |
| **Phloroglucinols** | 0.09 |
| **Pseudoalkaloids** | 0.05 |
| **Saccharides** | 0.17 |
| **Sesquiterpenoids** | 0.17 |
| **Small peptides** | 0.08 |
| **Steroids** | 0.07 |
| **Stilbenoids** | 0.12 |
| **Styrylpyrones** | 0.19 |
| **Triterpenoids** | 0.09 |
| **Tryptophan alkaloids** | 0.05 |
| **Tyrosine alkaloids** | 0.04 |
| **Xanthones** | 0.01 |

Supplementary Table 5: *Spec2Class* results on external validation with different confidence levels, applied by p1-p2 threshold

| number of spectra in the external validation set that have passed the p1-p2 threshold | p1- p2 threshold | f1 score | precision | recall | balanced accuracy | accuracy |
|---|---|---|---|---|---|---|
| **332** | 0 | 0.673 | 0.714 | 0.672 | 0.379 | 0.672 |
| **308** | 0.1 | 0.702 | 0.745 | 0.701 | 0.419 | 0.701 |
| **297** | 0.2 | 0.722 | 0.765 | 0.724 | 0.441 | 0.724 |
| **287** | 0.3 | 0.732 | 0.772 | 0.735 | 0.452 | 0.735 |
| **275** | 0.4 | 0.750 | 0.805 | 0.749 | 0.491 | 0.749 |
| **257** | 0.5 | 0.795 | 0.846 | 0.790 | 0.544 | 0.790 |
| **239** | 0.6 | 0.832 | 0.880 | 0.824 | 0.598 | 0.824 |
| **215** | 0.7 | 0.855 | 0.882 | 0.851 | 0.609 | 0.851 |
| **197** | 0.8 | 0.873 | 0.903 | 0.868 | 0.673 | 0.868 |
| **156** | 0.9 | 0.885 | 0.904 | 0.885 | 0.691 | 0.885 |

Supplementary Table 6: Benchmarking of Spec2Class vs CANOPUS on external validation. Results are based on similar spectra with and without the application of p1-1p2 threshold

| **p1-p2 threshold** | **number of spectra in the external validation set that have passed the p1-p2 threshold and where predicted by *CANOPUS*** | ***Spec2Class* accuracy** | ***CANOPUS* accuracy** | ***Spec2Class* precision** | ***CANOPUS* precision** | ***Spec2Class* recall** | ***CANOPU S* recall** | ***Spec2Class* f1 score** | ***CANOPUS* f1 score** | ***Spec2Class* balanced accuracy** | ***CANOPUS* balanced accuracy** |
|---|---|---|---|---|---|---|---|---|---|---|---|
| 0 | 235 | 0.725 | 0.689 | 0.753 | 0.792 | 0.725 | 0.689 | 0.719 | 0.726 | 0.542 | 0.566 |
| 0.1 | 218 | 0.753 | 0.716 | 0.780 | 0.798 | 0.753 | 0.716 | 0.748 | 0.746 | 0.563 | 0.596 |
| 0.2 | 212 | 0.770 | 0.722 | 0.800 | 0.804 | 0.770 | 0.722 | 0.765 | 0.753 | 0.588 | 0.584 |
| 0.3 | 206 | 0.773 | 0.728 | 0.803 | 0.809 | 0.773 | 0.728 | 0.768 | 0.760 | 0.571 | 0.573 |
| 0.4 | 196 | 0.787 | 0.735 | 0.832 | 0.821 | 0.787 | 0.735 | 0.786 | 0.769 | 0.599 | 0.600 |
| 0.5 | 185 | 0.823 | 0.757 | 0.862 | 0.842 | 0.823 | 0.757 | 0.825 | 0.790 | 0.631 | 0.640 |
| 0.6 | 169 | 0.865 | 0.757 | 0.902 | 0.855 | 0.865 | 0.757 | 0.870 | 0.799 | 0.672 | 0.605 |
| 0.7 | 151 | 0.888 | 0.781 | 0.895 | 0.883 | 0.888 | 0.781 | 0.886 | 0.825 | 0.614 | 0.604 |
| 0.8 | 137 | 0.899 | 0.803 | 0.906 | 0.899 | 0.899 | 0.803 | 0.895 | 0.845 | 0.653 | 0.648 |
| 0.9 | 107 | 0.898 | 0.813 | 0.893 | 0.904 | 0.898 | 0.813 | 0.892 | 0.852 | 0.657 | 0.621 |

### Supplementary figures

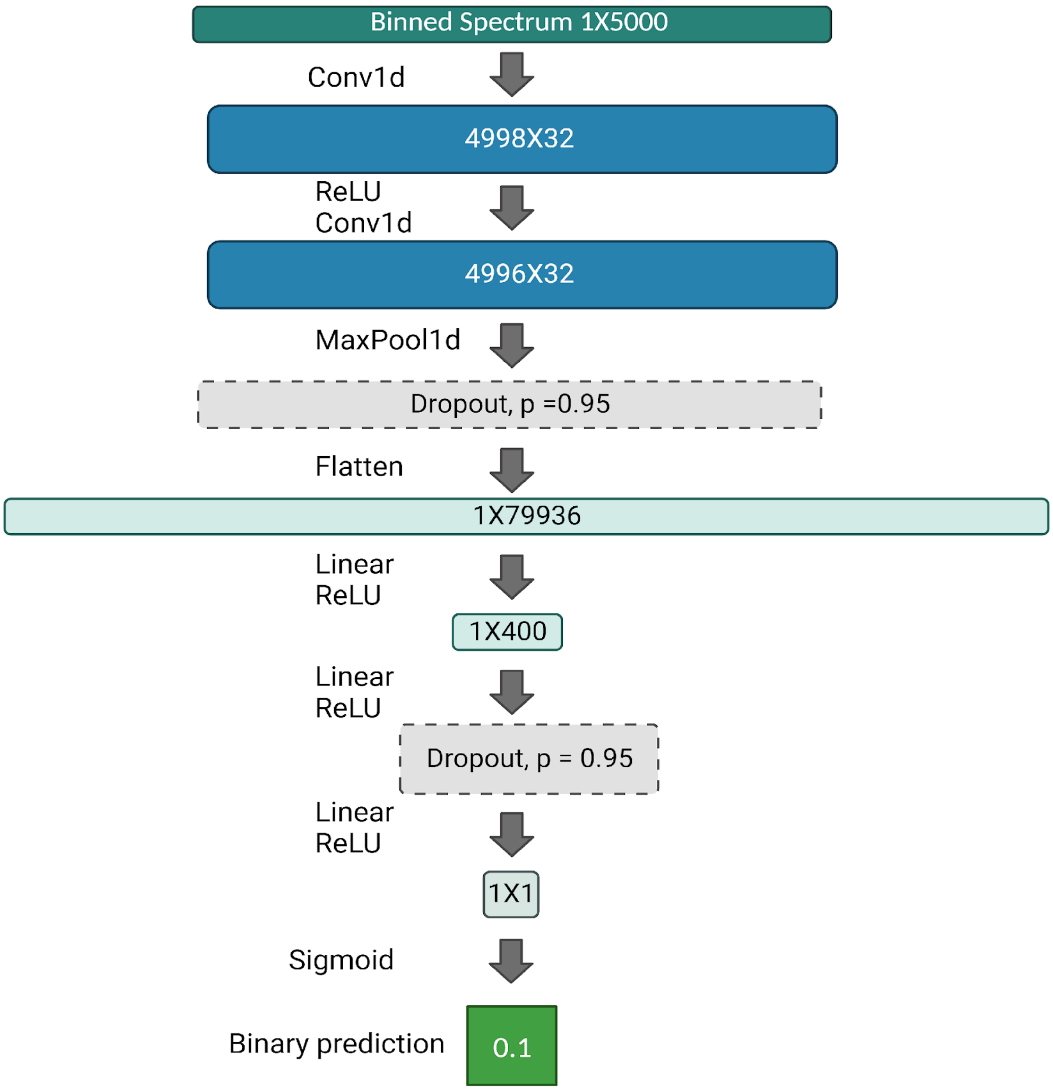


Supplementary Figure 1: The architecture of the neural net that was used as the chemical class binay model.

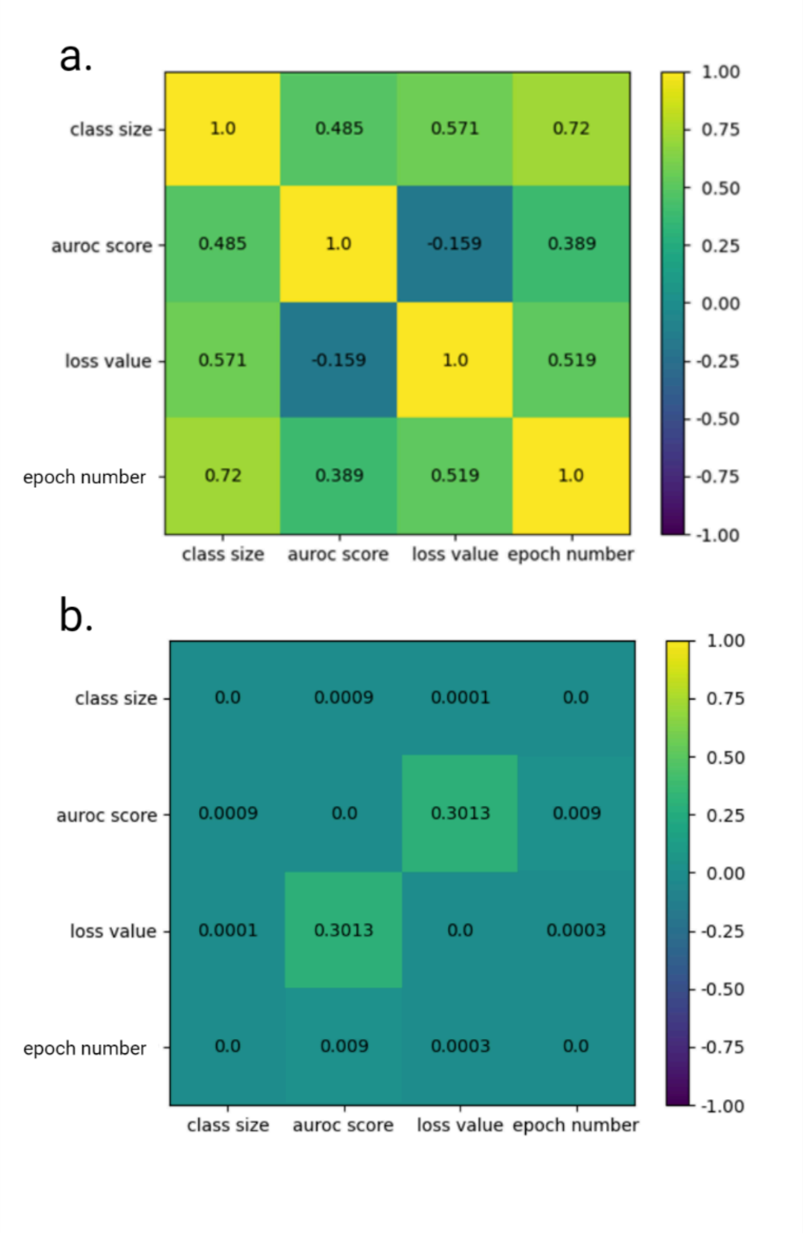


Supplementary Figure 2: Correlation and their significance based on binary model cross validation results. (a) Spearman correlation values between class size, auROC score and loss, based on cross validation results. (b) P value for the corresponding correlations in section

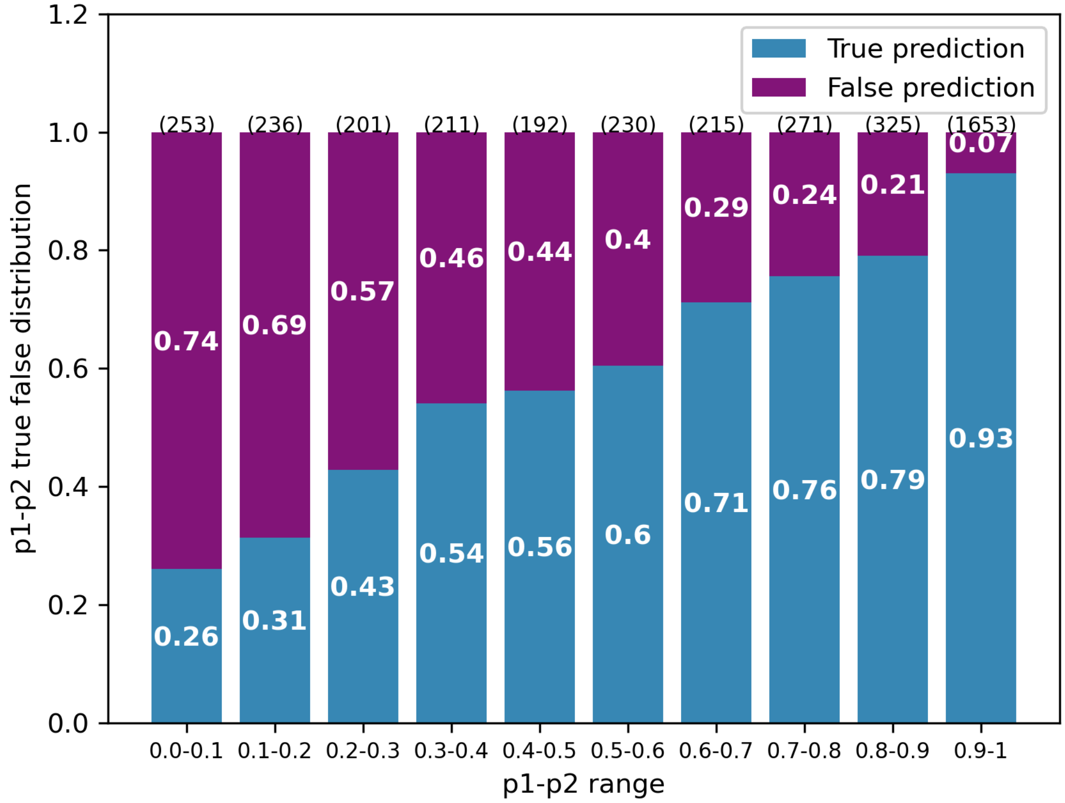


Supplementary Figure 3: The true and false prediction distribution for given p1-p2 values based on the test set results. For instance; a prediction with p1-p2 value that falls in the range 0.5-0.6, has the probability of 0.6 to be correct, based on 230 spectra from the test set.

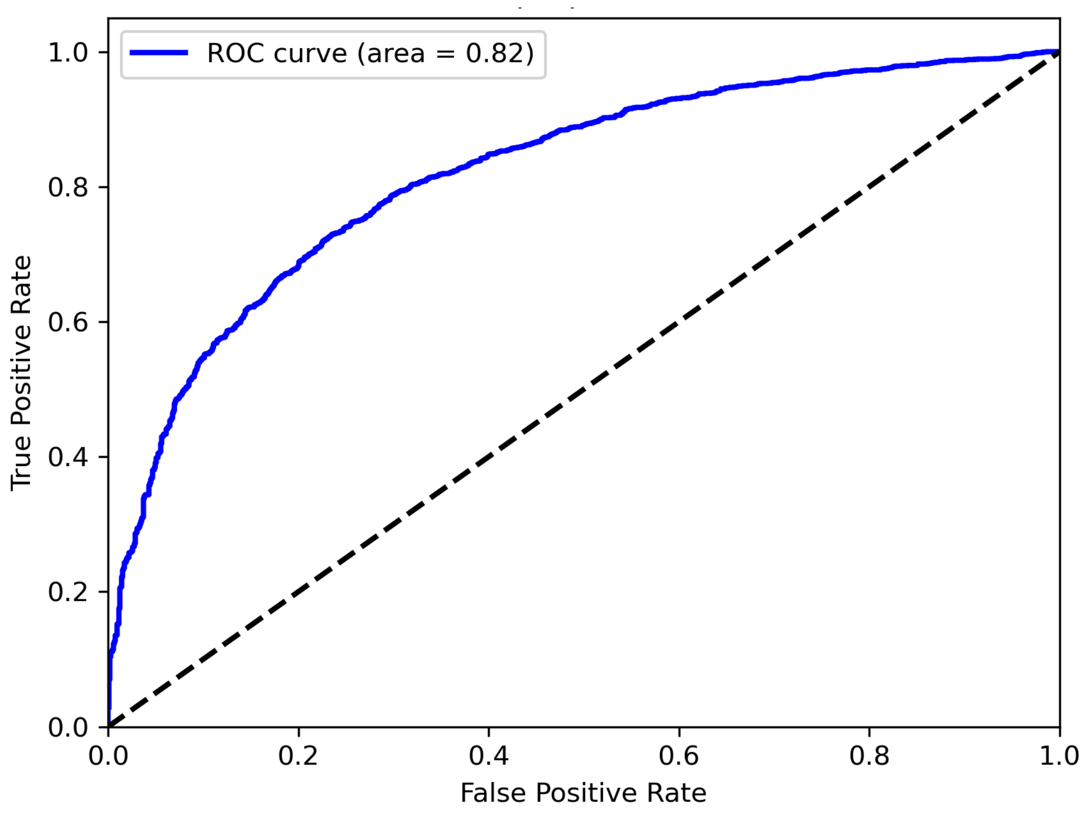


Supplementary Figure 4: Receiver operating characteristic (ROC) curve for p1-p2 different thresholds, based on the test set results. One can choose the p1-p2 threshold for the prediction analysis based on the True Positive, False Positive rates ratio (TPR/FPR). The high area under the ROC curve (auROC) value of 0.82 indicates the quality of the classification (true or false) based on the p1-p2 parameter.
